# An Improved 4’-Aminomethyltroxsalen-Based DNA Crosslinker for Biotinylation of DNA

**DOI:** 10.1101/2020.02.29.971317

**Authors:** Kevin Wielenberg, Miao Wang, Min Yang, Abdullah Ozer, John T. Lis, Hening Lin

## Abstract

Nucleic acid crosslinkers that covalently join commentary strands have applications as both pharmaceuticals and biochemical probes. Psoralen is a popular crosslinker moiety that reacts with double stranded DNA and RNA upon irradiation with long wave UV light. A commercially available compound EZ-Link Psoralen-PEG3-Biotin has been used in many studies to crosslink DNA and double strand RNA for genome-wide investigations. Here we present a novel probe, AP3B, which uses a psoralen derivative, 4’-aminomethyltrioxsalen, to biotinylate nucleic acids. We show that this compound is 8-fold more effective at labeling DNA in cells and several hundred-fold more effective at crosslinking two strands of DNA *in vitro* than the commercially available compound EZ-Link Psoralen-PEG3-Biotin.

## Introduction

Nucleic acid interstrand crosslinkers are important in nucleic acid research and come in many varities^1,2^. These range from simple molecules such as formaldehyde to inorganic complexes^3^ and can include polymers with a degree of specificity for a target DNA sequence^4^. Crosslinkers can covalently bond to nucleotides by reacting with nucleophilic regions of the bases or the double bonds of the bases. In the latter case activation of the compound by UV light is frequently required to induce pericyclic reactions^5,6^.

Photoactivable crosslinkers that can react with DNA and RNA in response to an external signal are particularly useful as they allow for spatial and temporal control.^7^ Psoralen is one such photoactivatable crosslinking moiety and has the added benefit of having natural affinity for DNA due to the compound’s planar multicyclic structure which allows it to intercalate nucleotide base pairs. Upon irradiation with UV light, psoralen undergoes 2+2 cycloaddition to pyrimidines; however, psoralen has specificity for reacting with thymines of complementary 5’-TA-3’ dinucleotide stretches^8^. This forms mono and bifunctional adducts,^9^ with bifunctional adducts crosslinking the two strands of the DNA (Figure 1).

**Figure 1.**
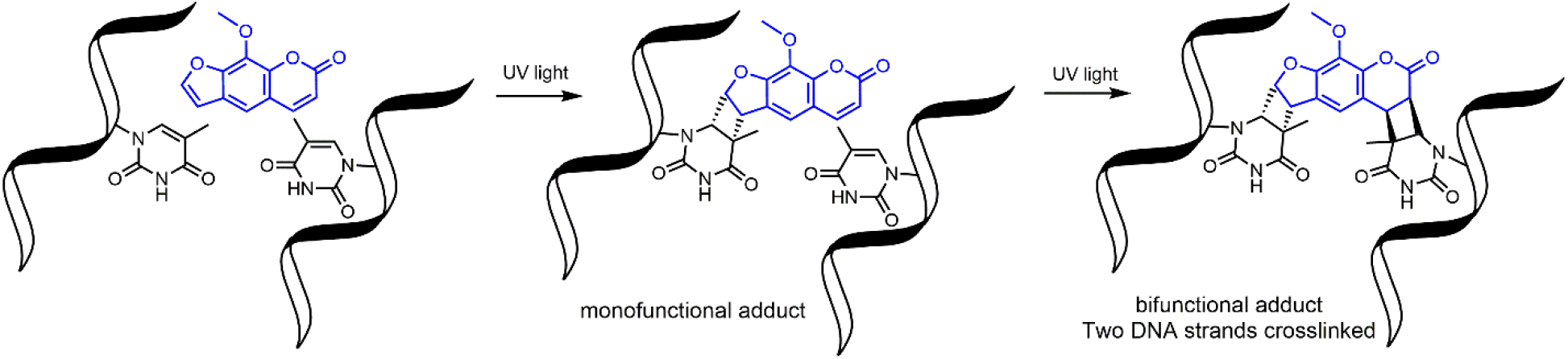
Psoralen undergo 2+2 cycloaddition reactions to form mono and bifunctional adducts with DNA.

EZ-Link Psoralen-PEG3-Biotin (PP3B, Figure 2A) is a commercially available probe that utilizes psoralen to biotinylate nucleic acids. Such crosslinkers enable cost effective biotinylation of DNA and RNA for further studies including sequencing of psoralen crosslinked, ligated and selected hybrids (SPLASH), studying RNA interactions, and Chem-seq to study the accessibility of DNA^10–13^. Despite the prevalent use of PP3B for these applications, we noticed that other psoralen derivatives such 4’-aminomethyltrioxsalen (AMT) are more effective nucleic acid crosslinkers than the psoralen methoxsalen derivative used in PP3B^14^. In this paper, we present the synthesis of a novel psoralen derivative-biotin compound, AMT-PEG3-biotin (AP3B, Figure 2B), and show that it is more efficient at labeling DNA than the commercially available PP3B.

**Figure 2.**
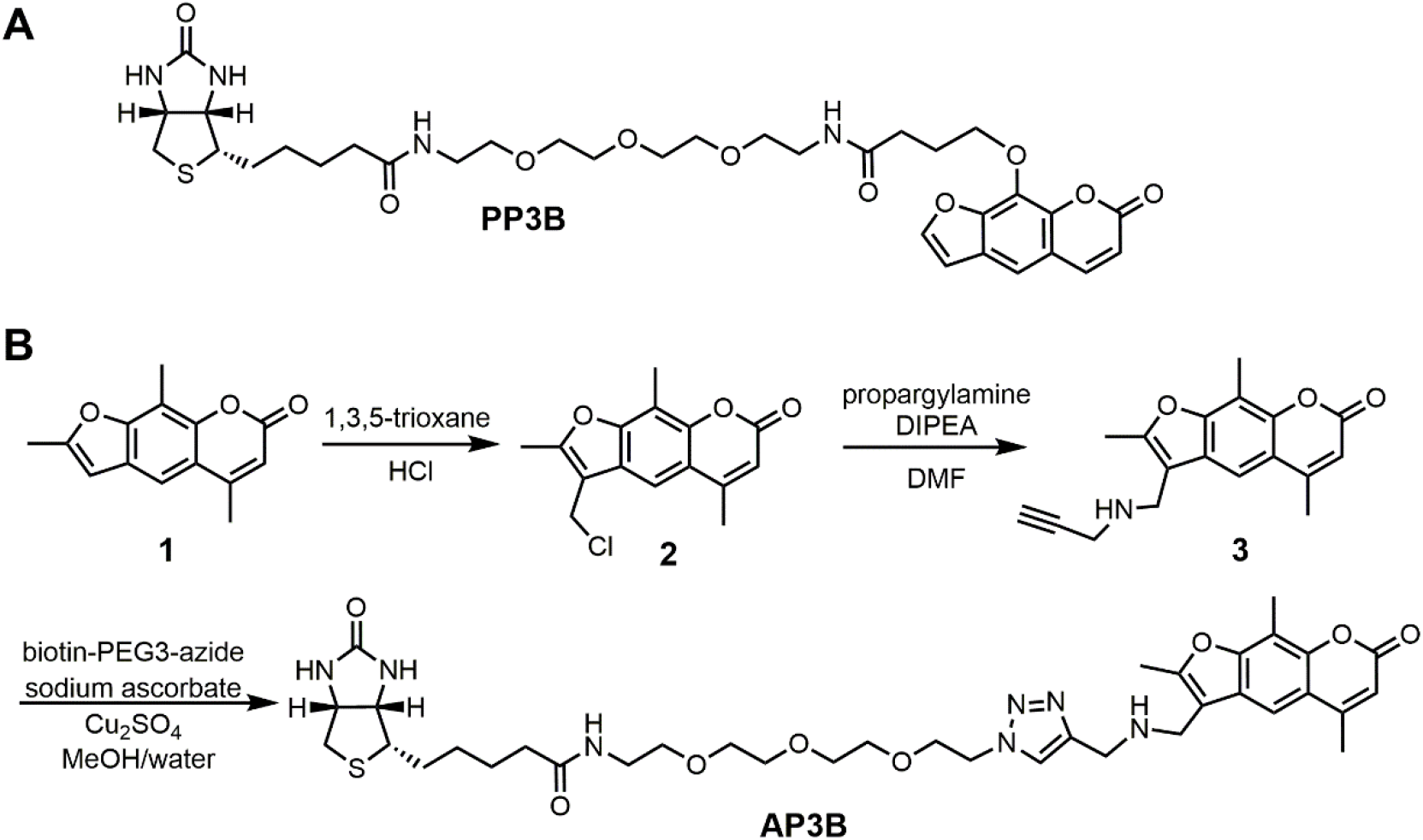
Structure of PP3B (**A**) and structure and synthesis of AP3B (**B**).

## Results and Discussion

### Design and Synthesis of AP3B

We decided to make a novel AMT containing nucleic acid probe as AMT has several advantages over psoralen and many other psoralen derivatives. AMT has increased affinity for DNA due in part to a positively charged amino group which is attracted to the negatively charged phosphate backbone of DNA and RNA. AMT also has better water solubility and a broader excitation spectrum which results in increased reactivity toward cycloaddition^15^. A PEG3 spacer was included to further increase water solubility. Biotin is used as the affinity tag since biotin streptavidin interaction is the best affinity reagent, with 10^-14^ M affinity and tolerance to harsh conditions such as high temperature (up to 65 °C) and denaturing conditions (up to 0.5% SDS). The resulting compound is AMT-PEG3-Biotin (AP3B, Figure 2B).

AP3B was synthesized from commercially available starting materials (Figure 2B). Briefly, 4’-chloromethyl-4,5’,8-trimethylpsoralen was reacted with 1,3,5 trioxane and concentrated HCl to attach a chloromethyl group to the furan ring. The alkyl chloride was then reacted with propargyl amine to attach a terminal alkyne which was then reacted with biotin PEG3 azide via click chemistry to produce AP3B.

### AP3B has increased crosslinking efficacy *in vitro*

We examined the efficiency of AP3B to crosslink purified DNA to that of the commercially available PP3B. For this purpose, we used an alkaline agarose gel mobility shift assay. Alkaline gels can disrupt base pairing in DNA to produce single stranded DNA. Crosslinking by AP3B or PP3B would prevent the two strands from separating producing up shifted bands. We first confirmed that AP3B crosslinking is dependent on UV irradiation (Figure 3A). We then compared the *in vitro* crosslinking efficacy with PP3B. The crosslinking band for 4 μM of AP3B had about the same intensity as 2,500 μM of PP3B (Figure 3B). Thus, AP3B is several hundred times more potent than PP3B *in vitro* at crosslinking the two strands of the DNA double helix.

**Figure 3.**
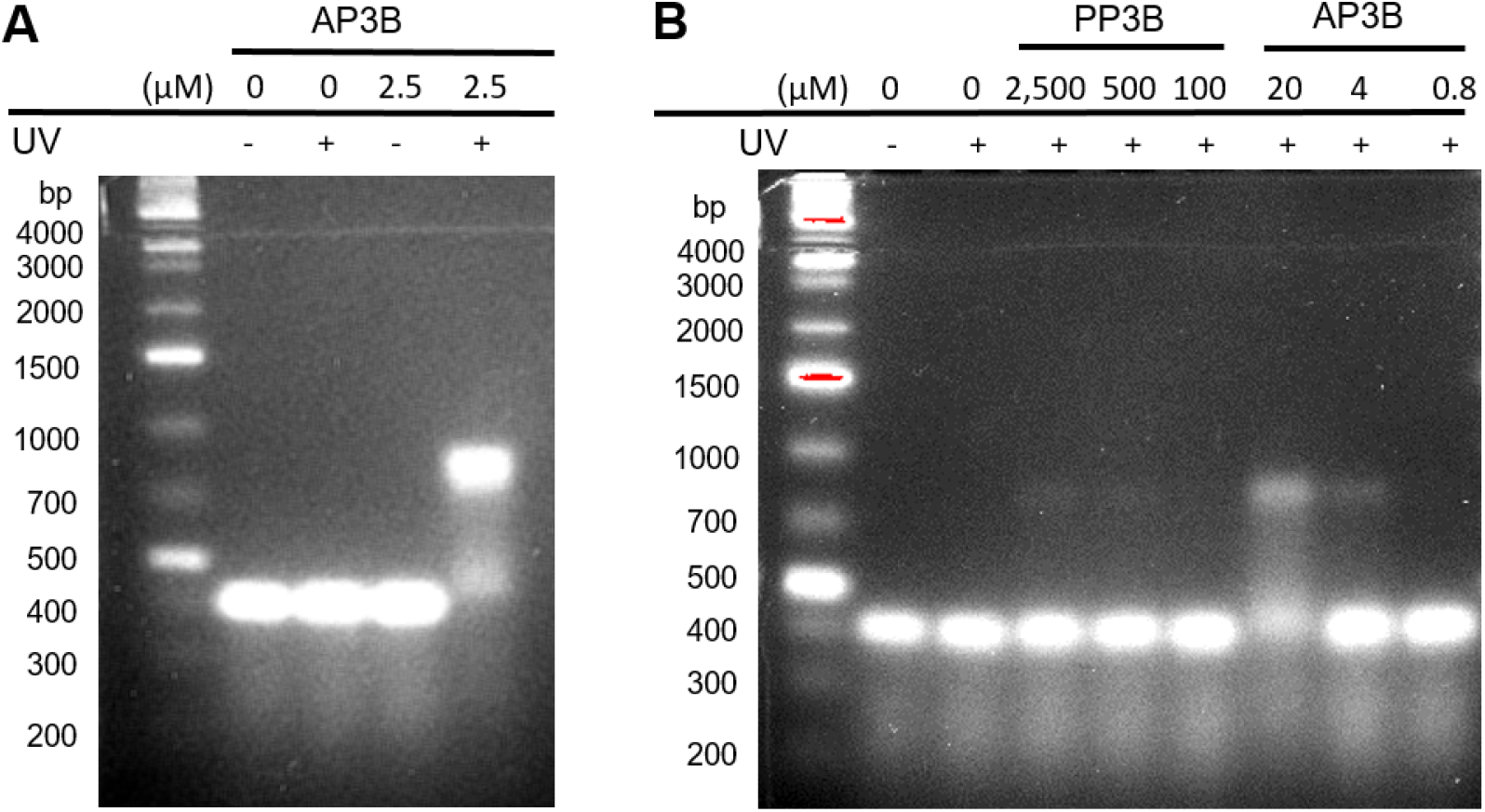
Detecting DNA crosslinking with alkaline agarose gels. These gels produce ssDNA by preventing base pairing between commentary strands. Crosslinking prevents separation of the strands resulting in a shift to higher molecular weight bands. (A) *AP3B* cross-linking shows dependence on UV irradiation. (B) Comparison of *PP3B* and *AP3B* crosslinking efficacy in vitro. AP3B shows significant signal depletion from the single stranded bands at concentrations where PP3B has no apparent crosslinking activity.

### AP3B shows higher labeling efficacy than PP3B

We next determined how the DNA labeling efficacy of AP3B compares to the commercially available PP3B which is commonly used in nucleic acid labeling experiments. An immunofluorescence assay was developed to compare the two compounds. Fixed cells were treated with PP3B or AP3B under UV light. Unreacted probes were washed away, and the covalently bound probes were recognized by a Streptavidin/rhodamine conjugate which was detected by florescence microscopy. Visual examination of the merged DAPI and rhodamine channels (Figure 4A) shows increased biotinylation in the AP3B treated cells at the same concentration as PP3B. This observation is confirmed by quantification of the signal intensity of rhodamine in cells’ nuclei (Figure 4B). AP3B’s labeling efficacy is over 3 times higher than PP3B at high concentrations and nearly 8-fold better at low concentrations.

**Figure 4.**
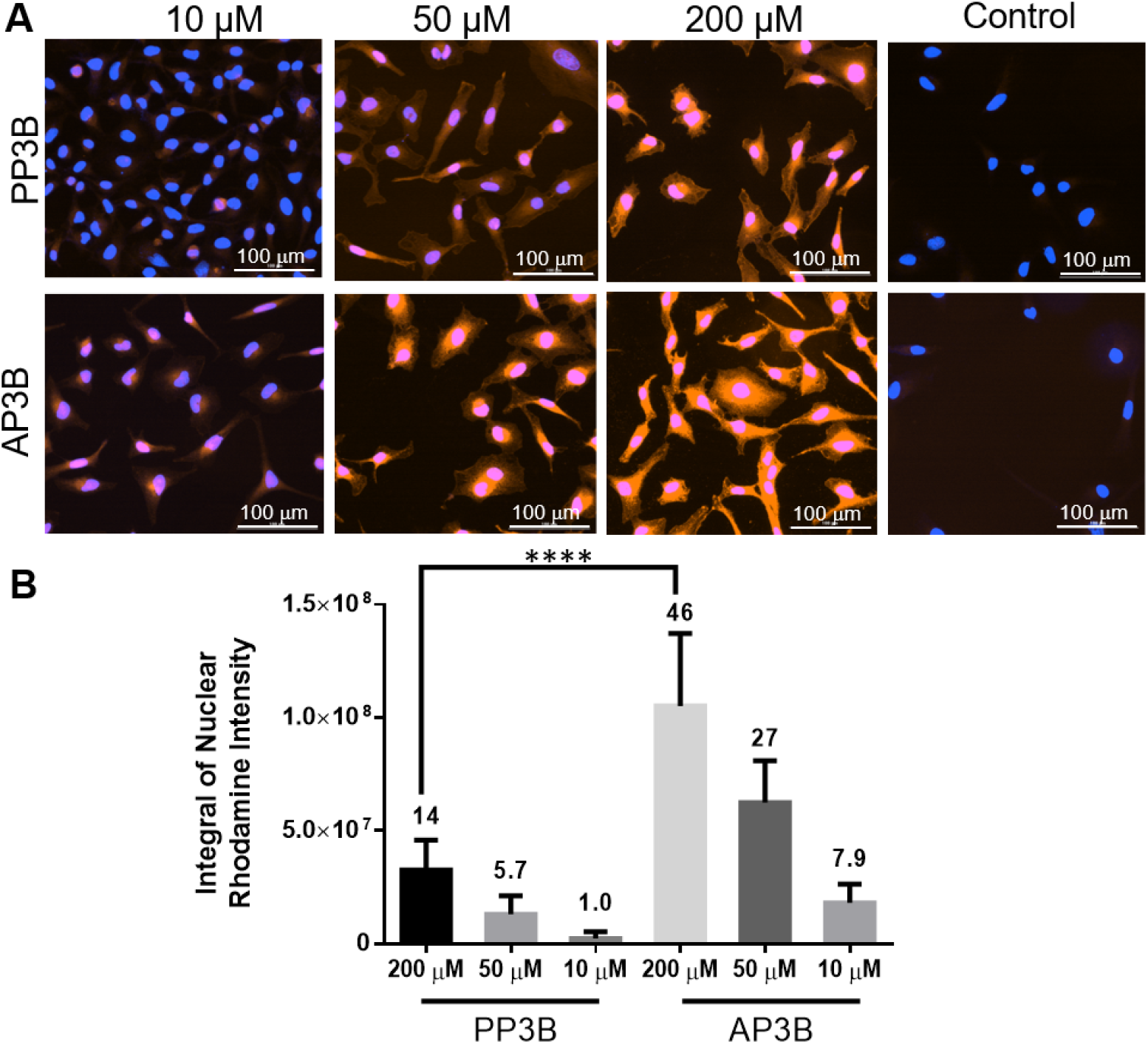
Evaluation of the relative labeling efficacy of PP3B and AP3B in cells. (A) In cell fluorescence detection of biotinylation in Hela cells incubated with the indicated concentration of PP3B or AP3B. (B) Quantification of the intergal of the nuclear rhodamine singal intensity with background (control) signal subtracted and bars normalized to the smallest integral.

In summary, we have synthesized a more efficient DNA crosslinking probe, AP3B containing a photoactivable AMT moiety. AMT is known to react more efficiently than psoralen and our experiments showed that AP3B is more reactive with DNA than the currently available product PP3B. Fluorescent detection of the compounds in cells shows an 8-fold increase in biotinylation for our compound compared to PP3B. This difference is further enhanced *in vitro* where AP3B is seen to produce hundreds of times more double strand crosslinks. The reason for the difference is likely because the in-cell fluorescence assay detects both mono and bifunctional adducts; however, the gel assay detects only bifunctional adducts that produce interstrand crosslinks between complementary strands of DNA. These results suggest that AP3B is a highly efficient tool for DNA biotinylation and could improve the efficacy of SPLASH^10^ and chem-seq^7^.

### Experimental Procedures

#### In-cell fluoresces assay

Hela cells were plated on 35 mm imaging petri dishes in DMEM supplemented with 10% FBS and grown for 24 h at 37 °C in an incubator with 5% CO_2_ air. The media was removed, and the cells were fixed with 4% paraformaldehyde for 15 min. Cells were washed three times with 1 mL of PBS to remove excess fixative. The cells were then incubated with 1 mL of blocking solution (1XPBS, 5% BSA, .1% Saponin) for 30 min at room temperature. Then PP3B or AP3B was added to the indicated concentration and incubated 10 min at room temperature. The solution was then removed, and the cells were washed with PBS to remove excess compound. The cells on the dishes were irradiated on ice with 368 nm light (5 1.2 W bulbs) at a distance of 6 cm for 30 min using a UV oven. The cells were then washed 3 times with blocking solution to remove unreacted molecules. The cells were then incubated with a streptavidin/rhodamine conjugate (TheremoFisher Scientific catalog number 21724) solution (1 μg/mL in blocking solution) at 4 °C for 1 h. Unbound streptavidin was removed with 3 washes with blocking solution (1 mL each). Anti-fade reagent containing DAPI (30 μL) was added to each dish and the fluorescent intensities of the DAPI and rhodamine signals were quantified using a Cytation 5 microscope.

#### DNA electrophoretic mobility shift assay

A denaturing gel (2% agarose) was prepared with alkaline buffer (30 mM NaOH, 2 mM EDTA). Solutions of purified DNA (30 ng/μL) and crosslinker at the indicated concentrations in PBS were irradiated on ice with a 368 nm light source (five 1.2 W bulbs) 6 cm away for 30 min using a UV oven. The samples and DNA ladders were then basified with 6x alkaline buffer and heat denatured at 70 °C for 5 min prior to loading onto the gel. The gel was then run for 3 h at 65 V. The gel was stained by soaking in 0.5 M Tris (pH 7.5) and SYBR Gold for 18 h prior to imaging.

#### Synthesis of 4’-Chloromethyl-4,5’,8-trimethylpsoralen (2)

To a solution of 1,3,5 Trioxane (0.71g, 7.9 mmol) in concentrated HCl (45 mL) was added **1** (2.05g 9.0 mmol). The reaction mixture was stirred at room temperature for 16 h. The reaction mixture was diluted with water (50 mL) and then extracted three times with chloroform (100 mL each). The organic layers were combined, washed once with brine (100 mL), and dried with Na_2_SO_4_ and then concentrated.

The crude product was recrystallized from ethyl acetate to afford the final compound (1.19g, 4.3 mmol, 54%). ^1^H NMR (500 MHz, Chloroform-*d*) δ 7.59 (s, 1H), 6.26 (s, 1H), 4.74 (s, *2*H), 2.57 (s, 3H), 2.52 (s, 6H); ^13^C NMR (126 MHz, Chloroform-*d*) δ 161.3, 155.2, 154.6, 153.12, 149.5, 123.9, 116.4, 113.2, 112.2, 111.2, 109.6, 36.2, 19.4, 12.3, 8.5.

#### Synthesis of AMT-alkyne (3). 2

(0.90g, 0.36 mmol) and propargylamine (0.23 mL, 36 mmol) were combined in toluene (50 mL). The reaction mixture was refluxed for 16 h after which the solution was cooled to room temperature and concentrated with a rotary evaporator. The crude residue was then purified by column chromatography (Hexane: EtOAc 1:1 → 1:2) to give the final product (0.80g 76%). ^1^H NMR (500 MHz, DMSO): δ 7.79 (s, 1H), 6.31 (s, 1H), 3.87 (s, 2H), 3.27 (s, 2H), 3.14 (s, 1H) 2.47 (s, 6H) 2.44 (s,3H); ^13^C NMR (126 MHz, DMSO) δ 160.1, 154.1, 153.9, 153.7, 148.3, 125.3, 115.5, 112.8, 112.5, 112.0, 107.5, 82.9, 73.8, 40.1 36.2, 18.7, 11.9, 8.2;

#### Synthesis of AP3B. 3

(0.16g, .54 mmol) and Biotin-PEG3-Azide (0.24g, 0.54 mmol) were dissolved in a solution of methanol and water (1:1, 14 mL). NaHCO_3_ (64 mg, 0.76 mmol), Cu_2_SO_4_ ·5H_2_O (14 mg, .054 mmol) and sodium ascorbate (21 mg, 0.11 mmol) were added to the reaction vessel. This solution was stirred at room temperature for 16h. The solution was extracted three times with DCM (14 mL). The pooled organic layers were washed once with brine (30 mL), dried with Na_2_SO_4_, then the solvent was removed with a rotary evaporator. The crude was purified by column chromatography (DCM: MeOH 10:1, 8:1, and then 6:1) to give the final product (0.37g, 0.50 mmol, 93%) 1H NMR (500 MHz, Methanol-d4) δ 7.98 (s, 1H), 7.82 (s, 1H), 6.28 – 6.22 (m, 1H), 4.60 (t, J = 5.0 Hz, 2H), 4.49 (dd, J = 7.9, 4.9 Hz, 1H), 4.29 (dd, J = 7.9, 4.5 Hz, 1H), 3.97 (s, 4H), 3.91 (t, J = 5.1 Hz, 2H), 3.65 – 3.58 (m, 4H), 3.58 – 3.52 (m, 4H), 3.49 (t, J = 5.5 Hz, 2H), 3.22 – 3.14 (m, 1H), 2.92 (dd, J = 12.8, 4.9 Hz, 1H), 2.70 (d, J = 12.8 Hz, 1H), 2.56 (s, 3H), 2.49 (d, J = 13.7 Hz, 6H), 2.18 (t, J = 7.4 Hz, 2H), 1.78 – 1.51 (m, 4H), 1.41 (p, J = 8.7, 8.1 Hz, 2H); ^13^C NMR (126 MHz, Methanol-*d*_4_) δ 174.6, 164.7, 162.1, 155.3, 155.2, 154.5, 148.7, 145.1, 125.4, 123.8, 115.9, 112.2, 112.0, 111.5, 108.2, 70.13, 70.08, 70.02, 69.8, 69.2, 69.0, 62.0, 60.2, 55.6, 50.0, 42.7, 40.7, 39.6, 38.9, 35.3, 28.3, 28.1, 25.4, 18.1, 10.8, 7.0. HRMS (ESI) m/z: calculated for C_36_H_49_O_8_N_7_S [M+H]^+^ : 740.3436, found 740.3413.

## Acknowledgments

The project described was supported by T32GM008500 from the National Institute of General Medical Sciences and the NIH common Fund 4D Nucleome Program U01HL 129958. The content is solely the responsibility of the authors and does not necessarily represent the official views of the National Institute of General Medical Sciences or the National Institutes of Health. This work made use of the Cornell NMR facility, which is supported, in part by, the NSF through MRI award CHE-153-1632.

## References

1 Sharma, E., Sterne-Weiler, T., O’Hanlon, D. & Blencowe, Benjamin J. Global Mapping of Human RNA-RNA Interactions. Molecular Cell 62, 618–626, doi:https://doi.org/10.1016/j.molcel.2016.04.030 (2016).

2 Lu, Z. et al. RNA Duplex Map in Living Cells Reveals Higher-Order Transcriptome Structure. Cell 165, 1267–1279, doi:10.1016/j.cell.2016.04.028 (2016).

3 Dasari, S. & Tchounwou, P. B. Cisplatin in cancer therapy: molecular mechanisms of action. Eur J Pharmacol 740, 364–378, doi:10.1016/j.ejphar.2014.07.025 (2014).

4 Baraldi, P. G. et al. Novel benzoyl nitrogen mustard derivatives of pyrazole analogues of distamycin A: synthesis and antileukemic activity. Bioorganic & Medicinal Chemistry 7, 251–262, doi:https://doi.org/10.1016/S0968-0896(98)00205-3 (1999).

5 Sakamoto, T., Tanaka, Y. & Fujimoto, K. DNA Photo-Cross-Linking Using 3-Cyanovinylcarbazole Modified Oligonucleotide with Threoninol Linker. Organic Letters 17, 936–939, doi:10.1021/acs.orglett.5b00035 (2015).

6 Song, C.-X. & He, C. Bioorthogonal labeling of 5-hydroxymethylcytosine in genomic DNA and diazirine-based DNA photo-cross-linking probes. Acc Chem Res 44, 709–717, doi:10.1021/ar2000502 (2011).

7 Dekker, J. et al. The 4D nucleome project. Nature 549, 219–226, doi:10.1038/nature23884 (2017).

8 Zhen, W. P., Buchardt, O., Nielsen, H. & Nielsen, P. E. Site specificity of psoralen-DNA interstrand cross-linking determined by nuclease Bal31 digestion. Biochemistry 25, 6598–6603, doi:10.1021/bi00369a039 (1986).

9 Couvé-Privat, S., Macé, G., Rosselli, F. & Saparbaev, M. K. Psoralen-induced DNA adducts are substrates for the base excision repair pathway in human cells. Nucleic Acids Res 35, 5672–5682, doi:10.1093/nar/gkm592 (2007).

10 Aw, Jong Ghut A. et al. In Vivo Mapping of Eukaryotic RNA Interactomes Reveals Principles of Higher-Order Organization and Regulation. Molecular Cell 62, 603–617, doi:https://doi.org/10.1016/j.molcel.2016.04.028 (2016).

11 Dadonaite, B. et al. The structure of the influenza A virus genome. Nature Microbiology 4, 1781–1789, doi:10.1038/s41564-019-0513-7 (2019).

12 Zhong, X. et al. The zinc-finger protein ZFYVE1 modulates TLR3-mediated signaling by facilitating TLR3 ligand binding. Cellular & Molecular Immunology, doi:10.1038/s41423-019-0265-6 (2019).

13 Anders, L. et al. Genome-wide localization of small molecules. Nat Biotechnol 32, 92–96, doi:10.1038/nbt.2776 (2014).

14 Isaacs, S. T., Shen, C.-K. J., Hearst, J. E. & Rapoport, H. Synthesis and characterization of new psoralen derivatives with superior photoreactivity with DNA and RNA. Biochemistry 16, 1058–1064, doi:10.1021/bi00625a005 (1977).

15 Oroskar, A., Olack, G., Peak, M. J. & Gasparro, F. P. 4’-Aminomethyl-4,5’,8-trimethylpsoralen photochemistry: the effect of concentration and UVA fluence on photoadduct formation in poly(dA-dT) and calf thymus DNA. Photochem Photobiol 60, 567–573, doi:10.1111/j.1751-1097.1994.tb05149.x (1994).

